# Growth- and stress-related defects associated to wall hypoacetylation are strigolactone-dependent

**DOI:** 10.1101/268417

**Authors:** Vicente Ramírez, Guangyan Xiong, Kiyoshi Mashiguchi, Shinjiro Yamaguchi, Markus Pauly

## Abstract

Mutants affected in the Arabidopsis TBL29/ESK1 xylan O-acetyltransferase display a strong reduction in total wall O-acetylation accompanied by a dwarfed plant stature, collapsed xylem morphology, and enhanced freezing tolerance. A newly identified *tbl29/esk1* suppressor mutation affects the biosynthesis of strigolactones (SL) due to the reduced expression of the *MAX4* gene. Genetic and biochemical evidence suggests that blocking the biosynthesis of SL is sufficient to recover all developmental and stress-related defects associated with the TBL29/ESK1 loss of function without affecting its direct effect - reduced wall O-acetylation. Altered levels of the *MAX4* SL biosynthetic gene, reduced branch number, and higher levels of methyl carlactonoate, an active SL, were also found in *tbl29/esk1* plants consistent with a constitutive activation of the SL pathway. These results indicate that the reduction of O-acetyl substituents in xylan is not directly responsible for the observed *tbl29/esk1* phenotypes. Alternatively, plants may perceive defects in the structure of wall polymers and/or wall architecture activating the SL hormonal pathway as a compensatory mechanism.

## INTRODUCTION

The success of higher plants to colonize terrestrial habitats rests greatly upon the evolution of the vascular system, the xylem, a highly specialized tissue for conducting water, minerals, and nutrients from soil to leaves over long distances (Lucas et al., 2013). Xylem tissue is characterized by the presence of specialized cells with tubular-shape structures formed by reinforcement of their cell walls, followed by a controlled cell death allowing a continuous column of water (Greenberg, 1996; Fukuda, 1997). During differentiation, these xylem cells modify the composition and structure of their secondary wall to increase the mechanical strength preventing the collapse of the vascular system under the pressure of water transpiration and extensive upright growth (reviewed in Schuetz et al., 2012). So far, forward and reverse genetic approaches have indicated that a correct synthesis and/or coordinate deposition of the secondary wall components may be necessary for the proper formation of xylem cells and the functionality of the plant vascular system. Several *irregular xylem* (*irx)* mutants have been identified in multiple plant species including poplar (Xi et al., 2016), rice (Chiniquy et al., 2013; Gao et al., 2017), maize (Sindhu et al., 2007; Vermerris et al., 2010) and Arabidopsis (Jones et al., 2001; Taylor et al., 2003; Scheller and Ulvskov, 2010; Xiong et al., 2013; Pauly et al., 2013). Most of the *irx* mutants display structural alterations in the main polymer components of xylem secondary walls (i.e., cellulose, hemicelluloses, and/or lignin) and a concomitant collapse of the xylem cells resulting in a stunted growth phenotype and activation of stress responses.

The identification of three protein families involved in the *O*-acetylation of hemicellulosic polysaccharides [Reduced Wall Acetylation (RWA), Trichome Birefringence-Like (TBL) and Altered Xyloglucan 9 (AXY9)] brought to light the crucial function of xylan *O*-acetyl substituents in xylem development as the corresponding mutants with reduced xylan acetylation displayed also collapsed xylem morphology, dwarfism and stress-related phenotypes (Xiong et al., 2013; Yuan et al., 2016a, 2106b and 2016c; Manabe et al., 2013; Schultink et al., 2015). One of the most extensively studied mutants in this category is *tbl29/eskimo1*. Chemical analyses of mutant walls revealed that mutations in this gene caused a reduction in 2-O- and 3-O- monoacetylation of xylan:xylosyl-residues without causing apparent alterations in the xylan chain length and the abundance of the reducing end sequence. *tbl29* mutant plants show collapsed xylem and reduced plant growth correlated with a constitutive expression of stress-related genes and enhanced tolerance to salt stress and freezing (Xin and Browse, 1998; Xiong et al., 2013; Yuan et al., 2013). Replacing the acetyl-substitutents in the *tbl29* mutant with glucuronic-acid moieties through the expression of a glucuronic acid transferase lead to the abolishment of the irregular xylem phenotype and thus normal plant growth (Xiong et al., 2015; Mortimer et al., 2015). These data suggested that a properly acetylated xylan structure is required for proper xylem function.

However, in this study we describe a detailed characterization of a *tbl29* suppressor mutant identified through a genetic suppressor screen uncoupling reduced xylan acetylation from the irregular xylem phenotype and providing evidence of an hitherto unreported role of the phytohormone strigolactone (SL) in the perception of secondary cell wall defects.

## RESULTS

### Identification and characterization of Arabidopsis *tbl29* suppressors

A recessive mutation (*tbl29S*) was found to rescue *tbl29*-associated growth defects such as reduction in plant size, altered plant architecture and a dark-green appearance (Figure 1A-B). In order to identify the causative mutation we used bulk segregant analysis combined with whole genome sequencing. This approach allowed us to identify an insertion co-segregating with the *tbl29*-suppressed phenotype around position −1269 in the promoter region of the *MORE AXILLARY BRANCHES 4* (*MAX4*) gene. This insertion leads to a 95% reduction in *MAX4* gene expression in a *tbl29S* background compared to the *tbl29* parental line (Supp. Figure 1). To confirm that the reduced expression of *MAX4* is responsible for the suppression of the *tbl29* developmental phenotypes, we crossed *tbl29* with a previously characterized *max4-1* knockdown mutant (Sorefan et al, 2003). The *max4-1* mutation was also able to suppress the *tbl29* phenotypes (Figure 1), confirming that the *tbl29* suppression in *tbl29S* is due to a mutation in *MAX4*. Accordingly, we changed the name of *tbl29S* to an additional allele of *max4*, *max4-7*.

**Figure 1.**
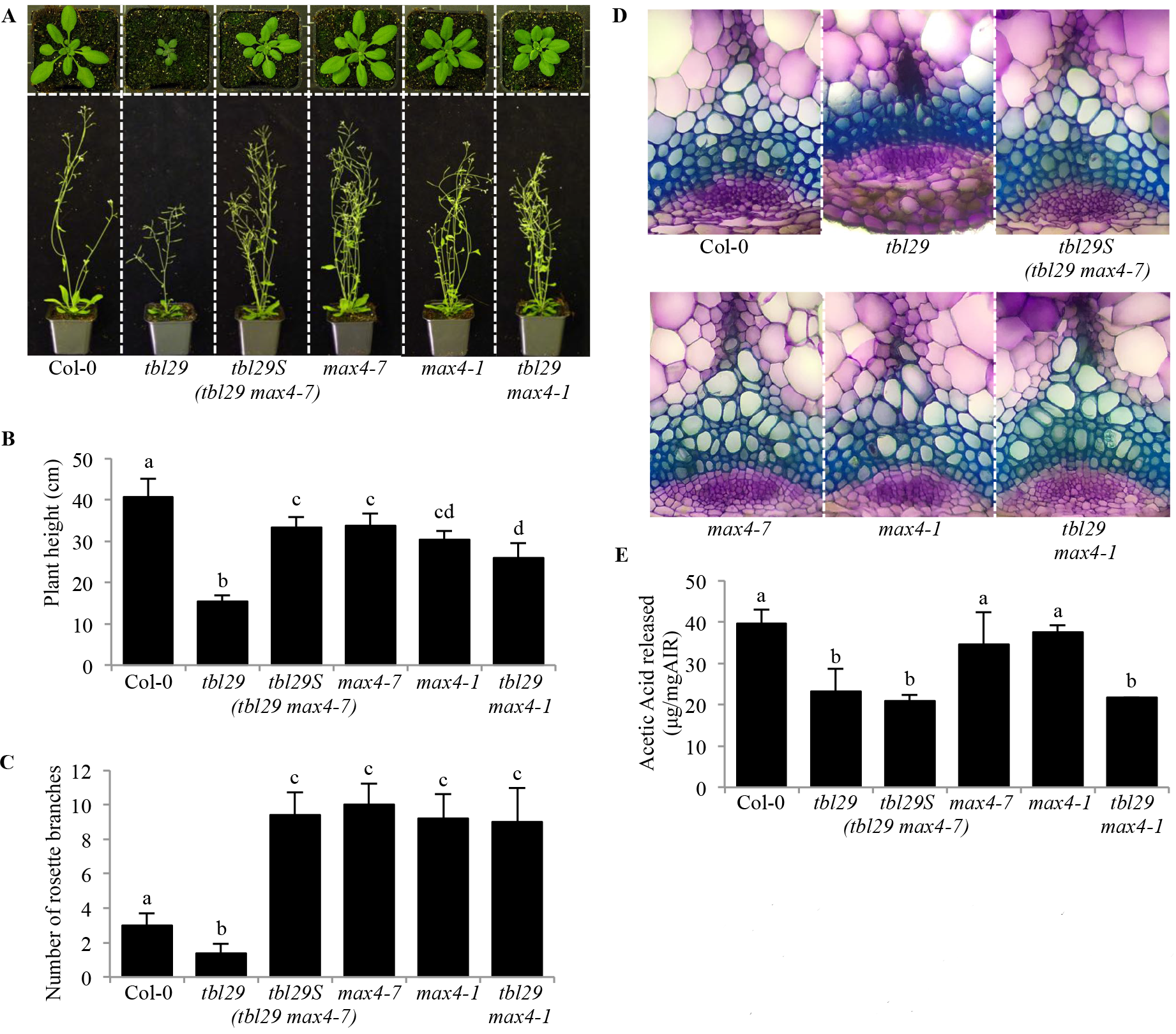
*tbl29S* mutation suppresses *tbl29* associated dwarfism, altered plant architecture and collapsed xylem. (**A**) Growth phenotypes of 4-week-old (upper panel) and 6-week-old (lower panel) plants. (**B**) Primary inflorescence heights (cm) (**C**) Number of rosette branches (**D**) Toluidine-O-Blue stained cross section of inflorescence stems. (**E**) Alkali-released acetate content of wall material (AIR) from stems. Data are represented as the mean of biological replicates (≥15 plants) with standard deviation. Means with different letters are significantly different (Tukev’s HSD, p<0.05)

A detailed characterization of the two suppressor alleles, *tbl29 max4-7* and *tbl29 max4-1*, was performed (Figure 1). Both double mutant plants lost the characteristic small size and dark-green color of the leaves (Figure 1A). The 62% reduction in stem height observed in the *tbl29* mutant is almost completely rescued in both *tbl29 max4-7* and *tbl29 max4-1* (Figure 1B). In addition, the altered shoot system architecture in *tbl29* is also rescued, measured as the rosette branching in mature plants (Figure 1C). *tbl29* plants have on average half the number of shoot branches arising from axillary meristems compared to Col-0. This reduced number of branches in *tbl29* is not only reverted to wildtype levels, but significantly increased in both double mutant alleles as previously described for other *max4* mutants (Sorefan et al, 2003).

Analysis of stem cross sections revealed that *max4* alleles also suppress the *tbl29*-associated irregular xylem morphology. In toluidine-blue stained sections from *tbl29* mutant plants, the vascular tissue contained irregular, collapsed-looking xylem elements. In contrast, in the equivalent stem sections from all plants containing *max4* mutations alone or in combination with *tbl29* showed a wildtype xylem morphology, where xylem elements can be easily identified as large round empty-looking cells with thick uniform cell walls (Figure 1D).

Cell wall compositional analyses on stem tissue showed that the reduced cellulose content in *tbl29* walls, which is typical among *irregular xylem* mutants, was fully restored in the *tbl29 max4-7* and *tbl29 max4-1*. The increased levels of non-cellulosic monosaccharides such as arabinose or xylose in *tbl29* stem tissue are also restored (Sup. Figure 2). In contrast, the characteristic low wall acetylation content in *tbl29* (42% reduction compared to wildtype) was retained in *tbl29 max4-7* and *tbl29 max4-1* plants (47% and 46% reduction respectively), implying that xylan acetylation remains low in the *tbl29*-suppressed plants. Taken together, these results suggest that blocking the expression of *MAX4* is able to suppress the characteristic dwarf phenotype and the collapsed xylem in a *tbl29* mutant background. Concomitantly, the indirectly altered cellulose and matrix polysaccharide levels and composition are restored. But this reversion does not involve a restoration of the direct effect of *TBL29* (xylan O-acetyltransferase) loss of function, reduced wall acetylation.

### Effect of exogenous SL application

MAX4 represents a carotenoid cleavage dioxygenase involved in the synthesis of SL (reviewed in Al-Babili and Bouwmeester, 2015). SL are a group of structurally similar terpenoid lactones controlling shoot branching. *max4* plants are affected in the production of carlactone (CL), an endogenous biosynthetic precursor of SL (Seto et al, 2014). Thus, mutant plants are highly branched, a phenotype that can be complemented by application of the synthetic SL GR24 (Gomez-Roldan et al., 2008; Umehara et al., 2008). To test if SL-deficiency was responsible for the suppression of *tbl29*-associated growth defects in the *tbl29 max4-7* double mutant, we examined the effect of exogenous applications of GR24 on plant architecture and cell wall composition of selected genotypes (Figure 2A-C). Supplementing a hydroponic culture media with 1μM GR24 did not result in any changes in WT plants (normal growth) or *tbl29* plants (dwarfed growth). However, when adding GR24 to the *tbl29 max4-7* double mutant, plant growth reverted back to dwarfism as found in the *tbl29* single mutant and the suppression through the *max4-7* mutation was abolished. Regarding plant architecture, GR24 treatment inhibited the axillary shoot growth in all genotypes tested except in *tbl29*, probably due to the already low branch number in this mutant under the growth conditions used (Figure 2C).

**Figure 2.**
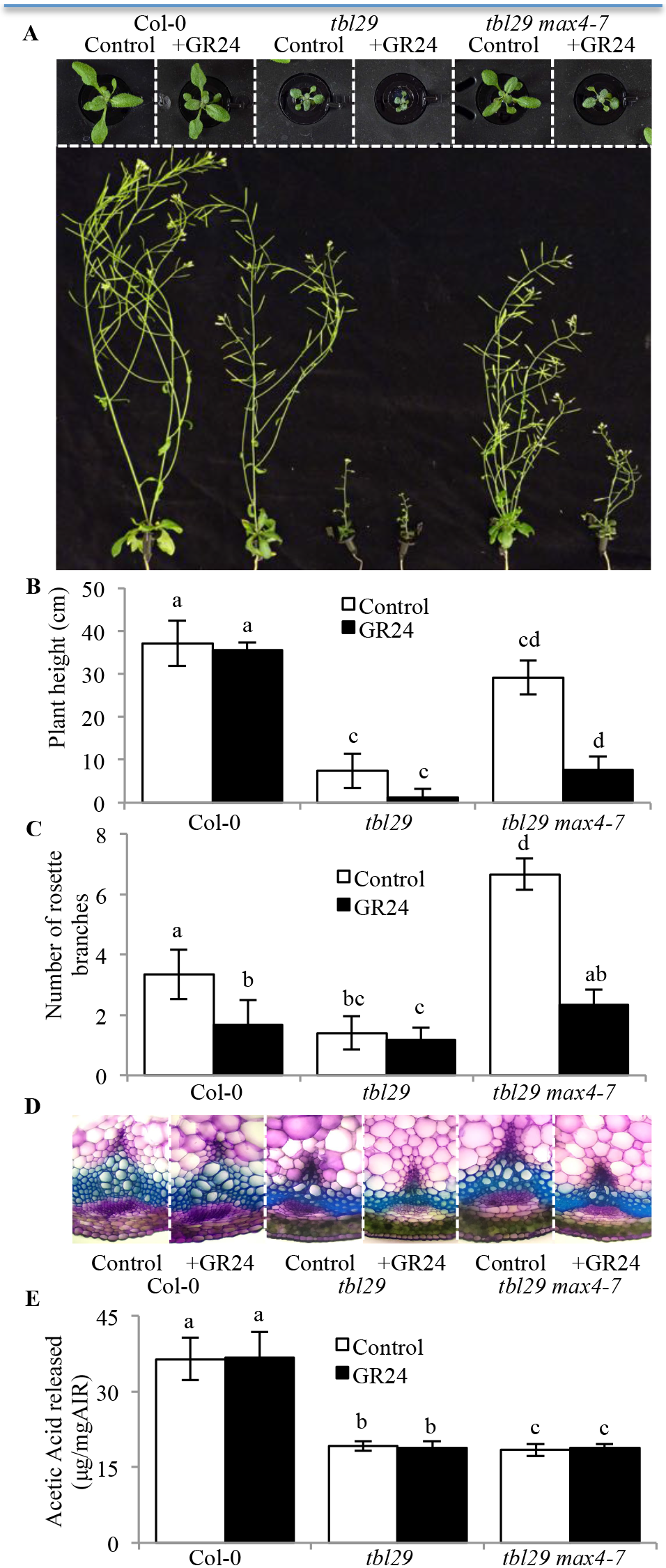
Effect of strigolactone application. (A) Growth phenotypes of 3-week-old (upper panel) and 6-week-old (lower panel) plants. (B) Primary inflorescence heights (cm) (C) Number of rosette branches (D) Toluidine-O-Blue stained cross section of inflorescence stems. (E) Alkali-released acetate content of wall material (AIR) from stems. Data are represented as the mean of biological replicates (≥5) with standard deviation. Means with different letters are significantly different (Tukey’s HSD, p<0.05)

Examinations of stems sections from GR24-supplemented *tbl29 max4-7* plants revealed collapsed xylem vessels similar to those characteristic of *tbl29* (Figure 2D).

Wall compositional analyses indicate that *tbl29 max4-7* plants treated with GR24 are similar to those observed for *tbl29* (Supp. Figure 3). In particular xylose levels increased in GR24-treated *tbl29 max4-7* plants to levels comparable to non-treated *tbl29* plants. No differences in the total wall acetate content in response to GR24 treatment were observed in any of the analyzed genotypes (Figure 2E).

Hence, the *max4-7* mutation is not able to complement *tbl29* in the presence of exogenously applied SL, indicating that a reduction in SL content suppresses the developmental defects including some wall structural defects caused by xylan hypoacetylation, but not hypoacetylation itself.

### Accumulation of active SL

Based on the results presented, a proper endogenous production of SL may be important for the expression of the highly pleiotropic phenotype of the *tbl29* mutant. In order to investigate if the reduced xylan acetylation in *tbl29* influences the synthesis of SL we measured the endogenous levels of methyl carlactonoate (MeCLA) in *tbl29* plants by LC-MS/MS. MeCLA is a non-canonical SL shown to be active in Arabidopsis plants (Abe et al., 2014). Endogenous MeCLA levels were not detectable in the two *max4* mutant alleles and the corresponding *tbl29 max4* double mutants confirming that *max4* mutations completely block MeCLA accumulation. The endogenous MeCLA content in Col-0 extracts was 103.3 ± 12.3 pg/g fresh weight and 156.47 ± 28.9 in *tbl29*, denoting a 50% steady state increase in the mutant (Figure 3). This result indicates that the biosynthesis of this SL is altered in *tbl29*.

**Figure 3.**
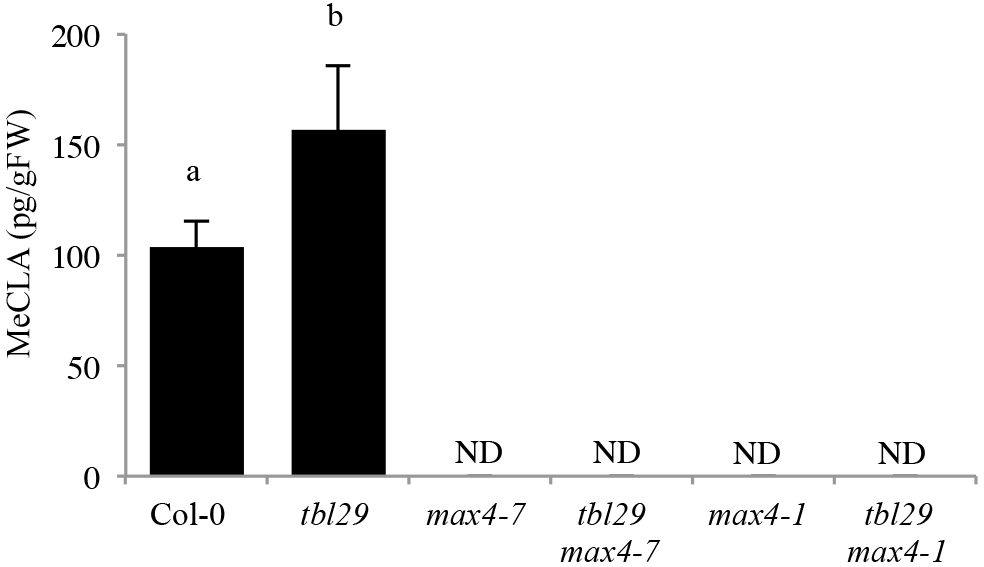
Quantification of endogenous MeCLA. Endogenous MeCLA was quan1fied by LC-MS/MS in Col-0, *tbl29*, *max4-7*, *tbl29 max4-7*, *max4-1* and *tbl29 max4-1* plants using D_1_-MeCLA as an internal standard. Data are represented as the mean of biological replicates (3-4) with standard devia1on. Means with different leLers are significantly different (Tukey’s HSD, p<0.05). FW, fresh weight. ND, not detectable.

### Suppression of *tbl29*-associated freezing tolerance

*tbl29* was first identified as *eskimo 1* (*esk1*) exhibiting an increased resistance to freezing temperatures (Xin and Browse, 1998; Xin et al., 2007). The survival of plants when subjected to freezing temperatures (−5°C) during a 20 hour period was tested. Under these conditions wildtype plants were reproducibly killed while most of the *tbl29* plants did not show any visible injury and were capable of continued growth after freezing and completion of their life cycle. To obtain a quantitative measure of freezing tolerance in the different genotypes, we determined the percentage of plants that are alive after the freezing treatment (Figure 4). The survival rate of *tbl29* in these freezing assays exceeded 90%, in contrast to Col-0 (3.3%). Interestingly, *tbl29 max4-7* plants showed the lethal wildtype-like phenotype with a survival rate of only 10%. These results indicate that blocking SL synthesis also suppresses the *tbl29/esk1*-associated freezing tolerance.

**Figure 4.**
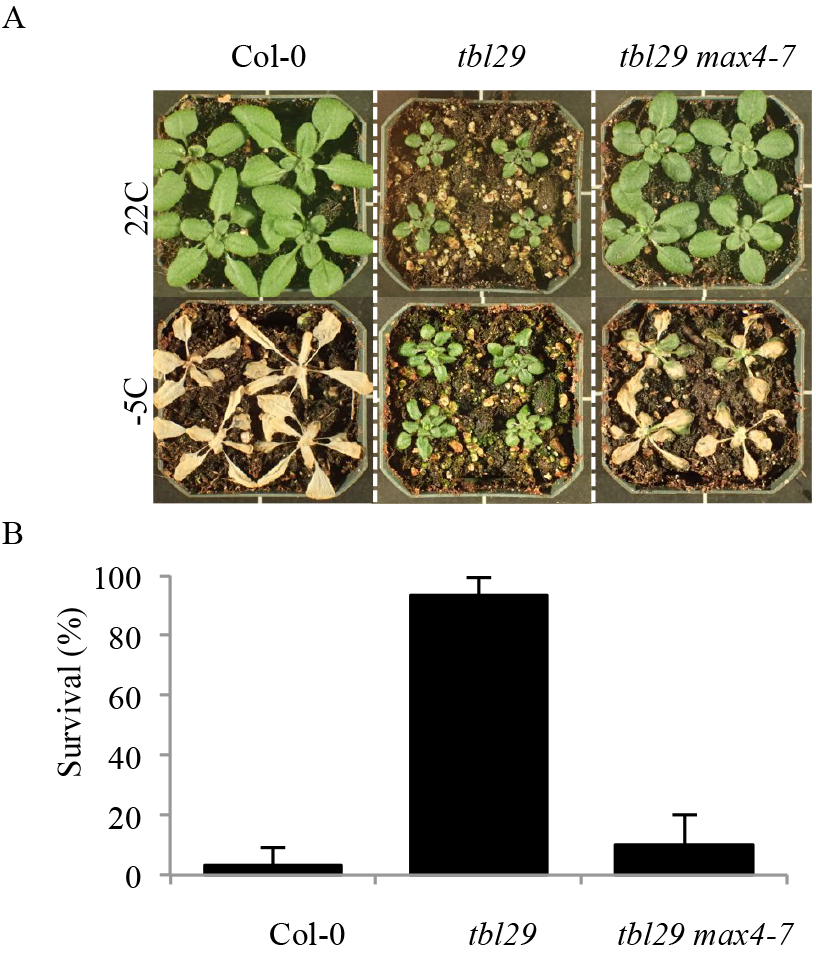
*tbl29*-associated freezing tolerance is suppressed by max4-7. (A) 4-week-old plants before (upper panel) and 3 days after freezing treatment (lower panel) Survival rate calculated as the percentage of plants alive after treatment. Error bars represent standard deviation (N=3 experiments, 20 plants per experiment)

## DISCUSSION

### Reduced wall acetylation can be uncoupled from dwarfism and collapsed xylem phenotypes

The functionality of the plant vascular system is dependent on the correct deposition and reinforcement of a secondary cell wall in xylem cells allowing the plant to resist the negative forces generated during water transport. The physiochemical properties of the secondary wall depend on the structure of the polymers (mainly cellulose, xylan and lignin) forming these walls. The degree of O-acetylation of xylan, the major acetylated polymer in secondary walls, seems to have a strong impact on these properties affecting the xylan-cellulose interaction (Ebringerová et al., 2005; Grantham et al., 2017). Although the biological significance of this modification is not clear, single and multiple mutants characterized in several plant species have shown a strong correlation between the level of reduction in xylan acetylation and a stunted plant growth and xylem collapse. The strongest reduction in wall O-acetylation has been reported so far for the *Arabidopsis axy9* mutant, with a 70% overall reduction affecting multiple polymers including xylan and another hemicellulose, xyloglucan (Schultink et al., 2015). No growth phenotypes have been reported for mutants lacking xyloglucan O-acetylation, so the severe growth defects, including dwarfed organs, a dark green leaf color and an extreme collapsed xylem, exhibited by the mutant has been attributed to the 80% reduction of xylan O-acetylation in stem tissue of *axy9*. The *Arabidopsis rwa2* single mutant exhibits a moderate 20% reduction in overall wall O-acetylation with no apparent plant development and/or xylem morphology defects, but an additive effect was observed by simultaneous mutation of multiple *RWA* genes (Lee et al., 2011). The quadruple *rwa* mutant shows a 42% reduction in xylan acetylation associated with a reduction in secondary wall thickening and collapsed xylem morphology. In the case of TBL family, several genes have been involved in xylan acetylation, with additive degrees of impact on O-acetylation and developmental defects. In rice, the *OsTBL1* and *OsTBL2* genes have recently been shown to be involved in xylan monoacetylation. The reduced growth and collapsed xylem in the double mutant is accompanied by a 55% reduction on total wall acetylation (Gao et al., 2017). In *Arabidopsis*, mutants affecting *TBL3*, *TBL31*, *TBL32*, *TBL33*, *TBL34* and *TBL35* genes show a minor decrease on xylan acetylation when analyzed individually but several double mutant combinations have been reported to cause moderate reductions ranging from 7% to 20% (Yuan et al., 2016a, 2016b, 2016c). Although a side-by-side comparison of all different mutant combinations has not been reported, there seems to be a direct correlation between the reduction of xylan acetylation and growth/ irregular xylem phenotypes. On the other hand, single loss-of-function alleles of the TBL29/ESK1 xylan acetyltransferase show a strong reduction on xylan O-acetylation (~ 40% depending on the report) accompanied by a strong arrest in plant growth and collapse of the xylem vessels (Figure 1; Supp. Fig. 2; Xin and Browse, 1998; Xin et al., 2007; Yuan et al., 2013; Xiong et al., 2013). *tbl29* mutant plants exhibit a reduction on the rosette size and overall plant height and a characteristic dark green color of the leaves. Interestingly, these growth defects can be complemented by replacing the missing acetyl-substitutents in the *tbl29* mutant with functionally equivalent glucuronic acid moieties through the expression of the AtGUX1 glycosyltransferase in the vascular tissue, suggesting that appropriately substituted xylan is required for proper xylem function (Xiong et al., 2015; Mortimer et al., 2010). However, we found a suppressor mutation able to complement all these *tbl29*-associated defects without affecting xylan hypoacetylation (Figure 1). This demonstrates that in *tbl29* reduced xylan O-acetylation is not directly responsible for the collapse of the xylem vessels and the corresponding developmental alterations. The structural defect of xylan is hence not essential for the xylem functionality and normal development. This notion of uncoupling xylan hypoacetylation and growth/ morphological defects is also supported by a previously identified *tbl29/esk1* suppressor mutation, *kaktus*, which was found to partially complement dwarfism and collapsed xylem also without recovering wall acetylation or other wall modifications (Bensussan et al., 2015). The *KAKTUS* (*KAK*) gene encodes a protein belonging to the E3-ubiquitin protein ligase family involved in endoreduplication and trichome architecture (Downes et al., 2003; El Refy et al., 2003). It is thought that *kak* is able to partially suppress *tbl29* dwarfism and collapsed xylem due to a hyperactivation of vascular development. However, the mechanism is unknown (Bensussan et al., 2015). A possible relationship between *KAK1* and SL requires further investigations.

### Irregular xylem and constitutive freezing tolerance phenotypes associated to wall hypoacetylation are SL-dependent

The *tbl29* suppressor mutation identified here is a new allele of *max4* (*max4-7*) representing an insertion in the promoter region of *MAX4*, causing a 95% reduction of the expression of the gene (Supp. Figure 1). A similar *tbl29* suppression effect was observed when crossing the mutant to a previously described *max4-1* allele, where a T-DNA insertion in the 1^st^ intron significantly impairs the expression of the *MAX4* gene (Sorefan et al., 2003), further confirming that a deficient MAX4 gene expression is responsible for the suppression of *tbl29* phenotypes. MAX4/CCD8 is a carotenoid cleaving deoxygenase catalyzing the biosynthesis of carlactone, the precursor of biologically active SL such as MeCLA in *Arabidopsis*. Consequently, *max4* mutants show a drastic reduction in SL accumulation (Figure 3; Seto et al., 2014; Abe et al., 2014). Genetic evidence demonstrates that blocking the endogenous production of SL is sufficient to suppress all *tbl29*-associated phenotypes as in the *tbl29 max4* double mutants (Figure 1; Supp. Figure 2). Moreover, complementing SL levels by exogenous applications of GR24, a synthetic SL, block the suppression effect of the *max4-7* mutation. While GR24 has a weak effect on wildtype plants, *tbl29 max4-7* plants treated with GR24 mimic the dwarf phenotype, collapsed xylem and most of the wall alteration attributes of *tbl29* control plants (Figure 2; Supp. Figure 3), suggesting that the reduction in SL synthesis is responsible for the *tbl29* suppression in the *tbl29 max4-7* double mutant. How blocking the SL biosynthesis in a secondary cell wall deficient mutant recovers the aberrant xylem development and prevents the activation of a general stress response requires further investigation, but several lines of evidence indicate that the SL pathway might be involved. Multiple indications in this report suggest that *tbl29* plants have a conspicuous alteration in the SL biosynthesis/perception. First, *tbl29* plants present a reduced branching phenotype (Figure 1C), consistent with the main role of SL as repressors of the outgrowth of the axillary bud (Gomez-Roldan et al., 2008; Umehara et al., 2008). The branch number has been used as typical phenotypic effect of SL signaling. While SL-deficient or SL-insensitive mutants show highly branched phenotype, activation of SL pathway has been associated with a reduction in branching both endogenously, through nutrient deficiency-triggered SL activation, or exogenously upon application of GR24 (Ito et al., 2016; Mashiguchi et al., 2009). Second, *tbl29* shows a 50% increased accumulation of active SLs (i.e. MeCLA) suggesting that SL biosynthesis is altered (Figure 3). Third, *MAX4* gene expression is down-regulated in *tbl29* stems (Supp. Figure 1), mimicking what has been reported for GR24-treated plants and opposed to the *MAX4* up-regulation observed in SL-deficient mutants due to a negative feedback regulation of SL biosynthetic genes observed in several plant species such as *Arabidopsis*, pea or rice (Mashiguchi et al., 2009; Johnson et al., 2006; Arite et al., 2007). Finally, an altered SL homeostasis in the *tbl29* mutant could also explain its general abiotic stress tolerance.

SL are carotenoid-derived plant hormones controlling multiple processes involved in plant development and adaptation to environmental cues (reviewed in Waters et al., 2017). Although there is some controversy, it has been suggested that SL positively regulates plant’s adaptation to abiotic stresses such as drought or salinity. In several plant species such as *Arabidopsis*, *Lotus japonica* or corn, GR24 applications increase the tolerance to drought, while conversely SL-deficient mutants show an increased sensitivity (Van Ha et al., 2014; Liu et al., 2015; Davidson et al., 2015). It is therefore likely that an increased endogenous SL content or perception in *tbl29* plants is part of the increased tolerance to multiple abiotic factors such as freezing, drought or salt stress (Figure 4; Xin et al., 2007; Bouchabke-Coussa et al., 2008; Xu et al., 2014). Accordingly, in the *tbl29 max4-7* double mutant, the *tbl29*-associated freezing tolerance is completely abolished (Figure 4). reinforcing the hypothesis that proper SL biosynthesis is necessary for the freezing tolerance presented by *tbl29/esk1* plants.

### Relationship between SL and xylem development

TBL29/ESK1 was first proposed to be a specific regulator of freezing tolerance (Xin and Browse, 1998), but several subsequent studies have reported an increased tolerance of *tbl29/esk1* mutants to abiotic factors including drought, salinity, osmotic stress or dehydration as well as biotic factors, indicating a more general role. Several metabolic alterations have been used to explain the constitutive tolerance of the *tbl29/esk1* mutant plants to these stresses such as increased accumulation of osmo-protectant metabolites (e.g. proline or soluble sugars), up-regulation of stress-specific subset of genes, or constitutive activation of the abscisic acid (ABA) signaling pathway (Lugan et al., 2009; Escudero et al., 2017). Recent reports favor the hypothesis that all these are indirect effects of the alteration in the water transport so the increased stress tolerance shown by the *tbl29/esk1* mutants is likely a consequence of the irregular xylem phenotype displayed by these plants (Bouchabke-Coussa et al., 2008; Lugan et al., 2009; Lefebvre et al., 2011, Bensussan et al., 2015).

The results presented in this manuscript indicate that a proper endogenous SL production or sensitivity is necessary for the development of the irregular xylem morphology in this mutant. How SL might regulate xylem morphology is still unknown, but we propose a model in which a defect in xylan acetylation in the secondary wall is somehow perceived by the cell by a hitherto unknown mechanism triggering the activation of a SL-dependent response. In the *tbl29* mutant, the continuous perception of a defective wall would lead to a constitutive activation of this signaling, leading to xylem collapse with the corresponding growth/ morphological/ stress response defects (Figure 5).

**Figure 5.**
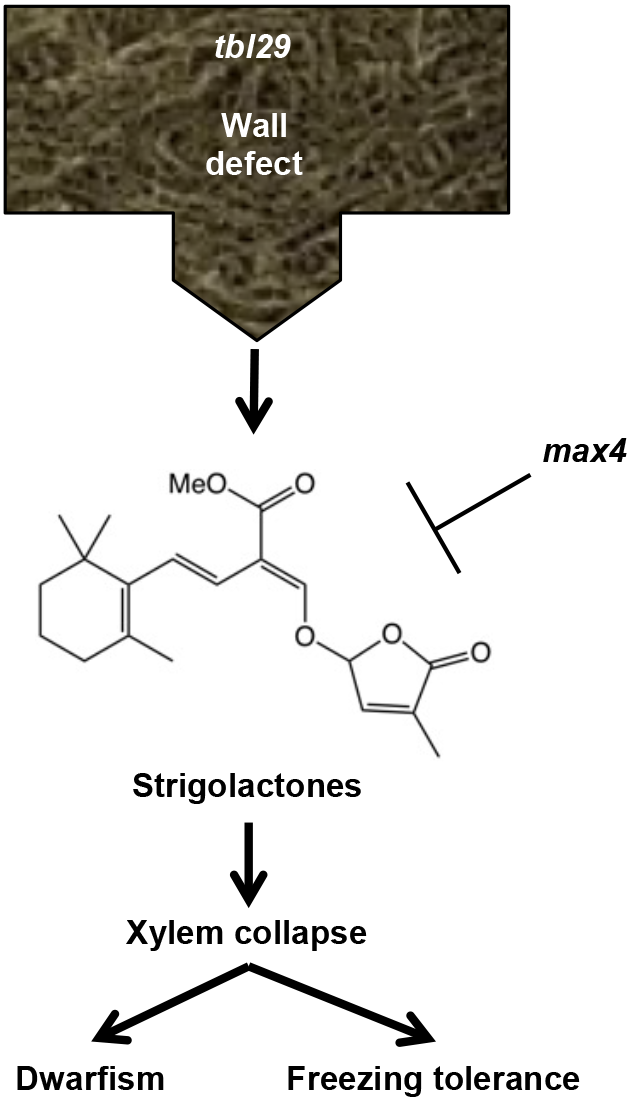
Proposed model. Defects in xylan acetylation in the secondary wall are perceived by an unknown mechanism triggering the activation of a SL-dependent response. In the *tbl29* mutant, the continuous perception of a defective wall would lead to a constitutive activation of this signaling, leading to xylem collapse with the corresponding growth/ morphological/ stress response defects. Altering the production of SL (as in the SL-deficient *max4* mutant) blocks the signaling preventing the developmental- and stress-related phenotypes.

Future work is needed to understand this SL-dependent cell wall sensing mechanism. The SL field is making great advances to identify and measure the SL compounds which are active in different plant species, and to unravel how they are perceived and the signal transduced in some SL-regulated processes (Reviewed in Lumba et al., 2017; and Waters et al., 2017). SL signal transduction controlling shoot branching seems to be based upon hormone-activated proteolysis of SL target proteins. There are a few SL targets proposed to be involved, including members of the SUPPRESSOR OF MAX2 1 (SMAX1) family (Stanga et al., 2013). The function of these proteins remains poorly understood, but some reports support the hypothesis that they are transcriptional regulators (Jiang et al., 2015; Soundappan et al., 2015). This is the case of other putative SL targets such as the BRI1-EMS SUPPRESSOR1 (BES1) and the DELLA family of GRAS transcriptional regulators, known targets of the brassinosteroid and gibberellin signaling, respectively (Nakamura et al., 2013; Wang et al., 2013). In that regard, some xylem-specific components of the wall biosynthesis/remodeling machinery are regulated by the SL pathway. For example, in hybrid aspen, a SL-dependent up-regulation of 1,3-β-glucanases (GH17 family, α-clade) controls shoot elongation via callose hydrolysis at sieve plates and plasmodesmata (Rinne et al., 2016). In addition, SL have been suggested as positive regulators of cambium activity in *Arabidopsis*, pea and in the woody plant *Eucalyptus globulus*. Although the molecular basis are unknown, local applications of GR24 in stems promote secondary growth by increasing the production of secondary vascular tissues and the lateral extension of the cambium zone (Agusti et al., 2011).

## MATERIALS AND METHODS

### Growth conditions

Unless otherwise indicated, seeds were stratified in sterile water for 4 days in the dark and 4°C, sown on soil and grown in environmentally controlled growth chambers under long day conditions (10000 luxes, 16h light, 22°C/8h dark, 19°C).

For the hydroponic experiment, seeds were surface-sterilized incubating them for 7 minutes in a 70% ethanol (Sigma-Aldrich), 0.1% Triton X-100 (Panreac Applichem) solution followed by five washes with sterile water. Seeds were then transferred to the Araponics seed holders (Araponics) filled with half-strength pH5.7 Murashige and Skoog media (Duchefa Biochemie) supplemented with 0.65% Plant Agar (Duchefa Biochemie) and incubated for four days in the dark at 4°C before being transferred to the growth chamber (7000 luxes, continuous day, 22°C). New Araponics nutrient solution (0.5mL/L) was added every week supplemented with rac-GR24 (Strigolab) or acetone as a control. After 6 weeks, plant height was recorded, pictures were taken, and stems were collected for toluidine-O-blue staining and cell wall analyses.

### Identification of *max4-7* (*tbl29S*)

A bulk segregant population was generated by crossing the *tbl29S* suppressor to the *tbl29* parental and 80 individuals in the F2 progeny were screened for presence/absence of the suppressed (dwarf) phenotype. Genomic DNA from the different individuals was prepared using the DNAeasy plant mini kit according to the manufacturer’s instructions (Qiagen) and equimolar amounts of DNA from the two groups were used for whole-genome sequencing. The 250PER Illumina sequencing was performed by the Vincent J. Coates Genomics Sequencing Laboratory in Berkeley using a Hi-Seq 2500 rapid instrument (Illumina). The obtained sequence reads were assembled using CLC Genomics Workbench 7.5 and aligned to the TAIR10 version of the Col-0 reference genome. Differences found in the suppressed and non-suppressed DNA pools were compared to identify the causative *tbl29S* mutation.

### Mutant genotyping

The different mutant lines were obtained from the NASC (http://arabidopsis.info) (Scholl et al., 2000) and ABRC (https://abrc.osu.edu) Arabidopsis stock centers. For the genotyping of the *tbl29* mutant (Stock number: SAIL_856_G11) and *max4-1* mutant (Stock number: CS9568), two PCR reactions were performed. Reaction A was performed using primers binding specific regions for each gene flanking the T-DNA insertion site (LP and RP primers). Reaction B was performed using a T-DNA specific primer (LB) and the RP primer used in reaction A. Similarly, the insertion in the *MAX4* (At4g32810) promoter region was confirmed using the LP and RP MAX4-specific primers flanking the insertion in reaction A, and in reaction B, the LP primer was combined with the insertion-specific primer LB. Primer sequences used for genotyping of the mutants described in this work are presented in Supp. Table 1.

### Plant measurements

The length of the primary steam (Plant height) and the number of shoot branches arising from axillary meristems (number of stems) were recorded 6 weeks after sowing the seeds and individual pictures were taken using a Lumix DMC-FZ35 camera (Panasonic).

### Xylem visualization

Plants were grown as indicated and the stems carefully collected. For comparative reasons, the middle of the first stem internode was selected for analysis. Hand-cut stem sections of the indicated genotypes were incubated one minute in 1mL 0.02% Toluidine Blue O solution (Sigma Aldrich) and then rinsed three times with 10mL sterile water. Sections were then gently pipetted onto a microscope slide, covered with a coverslip and observed under bright-field lighting microscope (Leica DM2000 LED).

### Cell wall composition

Individual primary stems were collected from 6-week-old plants, lyophilized using a ScanVac CoolSafe Freeze-dryer (Labogene) and homogenized to a fine powder using a MM400 mixer mill (Retsch Technology) at 30 Hz for two minutes. Total destarched alcohol insoluble residue (AIR) was extracted and one milligram per assay was used for the determination of cellulose content as previously described (Foster et al., 2010).

For the determination of matrix polysaccharide composition, 1mg of destarched cell wall material was hydrolyzed in 2M TFA for 90 min and 121°C and washed twice with 2-propanol before resuspension in H_2_O. A 940 Professional IC Vario HPAEC system with a pulse amperometric detector (Metrohm) was used to measure the different monosaccharides in the hydrolysate. Samples were separated with a Dionex CarboPac PA20 analytical column (3×150mm) and a Dionex CarboPac PA20 guard column (3×30mm) (Thermo Fischer Scientific), using a 0.4mL/min flow rate at 40°C. The eluents used were: A-Double distilled H_2_O; B-10mM NaOH (Sigma Aldrich); and C-700mM NaOH with the following elution profile: 20min 80%A+20%B, 5 min 30%A+70%C, 3min 100%C and 30 min 80%A+20%B.

Total wall acetate was determined using the 96-well-adapted format of the Acetic Acid Kit (Megazyme) as described previously (Guille et al., 2011).

### Analysis of methyl carlactonoate (MeCLA)

The analysis of MeCLA in Arabidopsis seedlings was performed as previously described (Abe et al., 2014). 13-day-old plants vertically grown on half strength Murashige and Skoog medium were homogenized in 5 mL of acetone containing [10-^2^H_1_-MeCLA (D_1_MeCLA) as an internal standard, by using POLYTORON PT3100D (Kinematica). The filtrates were evaporated under N_2_ gas and extracted twice with 4 ml of ethyl acetate after the addition of 3 ml of water and 1 ml of saturated sodium chloride solution. The organic phase was evaporated to dryness under N_2_ gas, dissolved in 0.4 ml of hexane, and loaded onto a Sep-Pak Silica cartridge column (1 cc; Waters). The column was washed with 2 ml of 5% (vol/vol) ethyl acetate/hexane and eluted with 2 ml of 15% (vol/vol) ethyl acetate/hexane. The eluate was then evaporated to dryness under N_2_ gas, dissolved in 0.3 ml of hexane, and loaded onto a Sep-Pak cyanopropyl cartridge column (1 cc; Waters), washed with 1 ml of hexane, and isocratically eluted with 3 ml of hexane. The eluate was evaporated to dryness, dissolved in acetonitrile, and then subjected to LC-MS/MS. LC-MS/MS analysis was carried out using a system consisting of TripleTOF 5600 (AB SCIEX) and Nexera ultra high performance liquid chromatography (Shimadzu) equipped with a CORTECS UPLC C18+ column (1.6 *μ*m, ϕ2.1 × 100 mm; Waters). Elution of the samples was carried out with 0.05% (vol/vol) acetic acid/water (solvent A) and 0.05% (vol/vol) acetic acid/acetonitrile (solvent B) using a gradient from 30% to 95% solvent B (0 min to 6.5 min) at a flow rate of 0.3 ml/min. The temperature of the column was set to 35°C. The MS/MS analysis was operated under the enhanced mass mode as following parameters: Declustering potential, 55 V; collision energy, 25 V; GS1, 35 psi; GS2, 35 psi; temperature, 400°C; precursor ion (*m/z*), 347.2 for MeCLA and 348.2 for D_1_-MeCLA. A fragment ion with *m/z* 97.03 was used to identify both MeCLA and D_1_-MeCLA. A standard curve for quantification was generated by using MeCLA and D_1_-MeCLA.

### Freezing tolerance assay

4-week-old plants grown in soil under the described conditions were incubated at −5°C in a Coolfreeze CF-110 cooling box (Waeco) for 20 hours and then transferred back to the growth chamber. After three days pictures were taken and the number of surviving plants recorded.

### Gene expression analysis

Primary stems from 6-week-old plants were collected and immediately frozen in liquid nitrogen. Tissue was homogenised to a fine powder using a MM400mixer mill (Retsch Technology) at 30 Hz for one minute and total RNA was extracted DNAse-treated using the Plant RNeasyMini Kit according to the manufacturer’s instructions (Qiagen). One microgram of RNA was then used for the synthesis of complementary DNA using the Tetro cDNA Synthesis Kit (Bioline) and one microliter was used as template to perform real-time quantitative PCR with SsoAdvanced Universal SYBR Green Super Mix (BioRad) in a CFX96 System (Biorad). Relative MAX4 transcript abundance was calculated using the ACTIN2 (At3g18780) gene as an internal control using the following primers: MAX4-F: 5’-TGTGGAGTAGCCGTCGAAGAG-3’, MAX4-R: 5’-GAAAGATACCCACTTGGCTGAATG-3’, ACT2-F: 5’-TCTTCCGCTCTTTCTTTCCAAGC-3’ and ACT2-R: 5’-ACCATTGTCACACACGATTGGTT G-3’

## ACKNOWLEDGEMENTS

We thank Ulrike Klauss for excellent technical assistance.

## ACCESSION NUMBERS

Sequence data from this article can be found in the EMBL/GenBank data libraries under accession numbers: *TBL29/ESK1*, At3g55990; *MAX4/CCD8*, At4g32810.

## SUPPLEMENTAL DATA

Supp. Figure 1. *tbl29S* mutation represents an insertion into the At4g32810 promoter region.

Supp. Figure 2. Monosaccharide composition, cellulose and acetate content of stem cell walls.

Supp. Figure 3. Monosaccharide composition, cellulose and acetate content of stem cell walls in control and GR24-treated plants growing in hydroponic culture.

Supp. Table 1. Primers used in this study.

